# Learning and memory in hybrid migratory songbirds - cognition as a reproductive isolating barrier across seasons

**DOI:** 10.1101/2023.06.07.544084

**Authors:** Ashley Alario, Marlene Trevino, Hannah Justen, Constance J. Woodman, Timothy C. Roth, Kira E. Delmore

## Abstract

Hybrid zones can be used to identify traits that maintain reproductive isolation and contribute to speciation. Cognitive traits may serve as post-mating reproductive isolating barriers, reducing the fitness of hybrids if, for example, misexpression occurs in hybrids and disrupts important neurological mechanisms. We tested this hypothesis in a hybrid zone between two subspecies of Swainson’s thrushes (*Catharus ustulatus*) using two cognitive tests – an associative learning spatial test and neophobia test. We included comparisons across the sexes and seasons (spring migration and winter), testing if hybrid females performed worse than males (as per Haldane’s rule) and if birds (regardless of ancestry or sex) performed better during migration, when they are building navigational maps and encountering new environments. We documented reduced cognitive abilities in hybrids, but this result was limited to males and winter. Hybrid females did not perform worse than males in either season. Although season was a significant predictor of performance, contrary to our prediction, all birds learned faster during the winter. The hypothesis that cognitive traits could serve as post-mating isolating barriers is relatively new; this is one of the first tests in a natural hybrid zone and non-food-caching species. We also provide one of the first comparisons of cognitive abilities between seasons. Future neurostructural and neurophysiological work should be used to examine mechanisms underlying our behavioral observations.

## Introduction

Hybrid zones are areas where closely related, but distinct, groups of populations meet, mate and produce offspring of mixed ancestry^[1,2]^. Many stable hybrid zones exist in nature^[3–6]^. The persistence of these hybrid zones suggests that some form of reproductive isolation exists between the groups; however, in many cases, we do not know which traits generate this isolation.

Cognitive traits including learning and memory may contribute to reproductive isolation in some systems. Specifically, reproductive isolation can be generated by both pre- and post-mating barriers^[7]^. Learning and memory likely contribute to pre-mating isolation as they are important for mate choice and can promote assortative mating (e.g., through sexual imprinting, with offspring using parental characteristics learned in life to select future mates^[8,9]^). These traits also affect individual-level fitness^[9–11]^ and can have a genetic basis^[12]^ suggesting learning and memory may also serve as post-mating barriers to gene flow.

Rice and McQuillan described two mechanisms by which learning and memory could serve as post-mating isolating barriers^[13]^. First, parental populations often inhabit different environments. Divergent selection on learning abilities in these environments could cause hybrids to exhibit intermediate abilities that do not match either parental environment. Second, if parental populations spent time in allopatry, alternate alleles at loci underlying cognitive traits could have fixed in each population and be incompatible in hybrids when they come back into contact. In some cases, these incompatibilities (whether they underlie cognitive traits or not) result in misexpression; mechanisms important for gene regulation diverge in allopatry and are lost when they recombine in hybrids causing genes to be over or under expressed compared to parental populations. Misexpression likely interferes with many traits in hybrids, including cognition^[13]^.

Very few studies have tested the role that cognitive traits could play as post-mating isolating barriers. Some work in lab-bred taxa suggests that hybrids may actually be more fit than parental forms (e.g., lab bred hybrid mules performed better in visual discrimination learning tasks) but, in many of these cases, this pattern may be related to inbreeding depression in parental forms^[14,15]^. Work in a natural hybrid zone between black-capped and Carolina chickadees has supported a role for cognition and speciation. Specifically, McQuillan et al. 2018 ^[16]^ compared the spatial learning and problem-solving abilities of these species and their hybrids. Chickadees are scatter hoarders that cache food for the winter^[17]^. Hybrids performed worse than their parental forms in both cognitive tasks, suggesting they may have lower fitness in the winter when they need to retrieve their food. Under this scenario, learning serves as a strong selection factor and hence a post-mating isolating barrier. Wagner et al.^[18]^ also documented indirect evidence of reduced introgression at genes important for neurological function in this hybrid zone. Additional research with other species and domains beyond food caching is required to test if cognitive traits play a more general role as post-mating isolating barriers. Here, we focus on a hybrid zone between Swainson’s thrushes (*Catharus ustulatus*), a non-food-caching bird, and focus on seasonal migratory behavior, comparing cognitive abilities across seasons and ancestry classes.

Seasonal migration is the yearly, long-distance movement of individuals between their breeding and wintering ranges. There is a strong genetic basis to this behavior in many groups (including songbirds), but learning is still important for successful migration and likely has important fitness consequences for migrants^[19,20]^. For example, many migrating species develop navigational maps from cues that they learn on the first leg of migration to help complete subsequent trips^[21]^. The development of these maps is beneficial, as they allow migrants to take more direct routes, revisit good stopover sites and avoid unfavorable areas^[22,23]^. In addition, migrants encounter new environments throughout their annual cycle, especially on migration when they are traveling over long distances, using stopover sites that can span different continents and climatic zones. Migrants only have a short amount of time to refuel at stopover sites and predation can be high^[24]^. Cognitive traits may help facilitate responses to new or changed environments (e.g., by allowing individuals to adopt new resources and/or avoid novel threats^[9,11]^) and could thus be important for ecological adaptation on migration.

The Swainson’s thrush (*Catharus ustulatus*) is a migratory songbird that includes two subspecies: coastal, russet-backed, and inland, olive-backed thrushes^[25]^. These subspecies form a narrow hybrid zone where their ranges overlap in western North America^[26]^ and migration is the main trait that distinguishes them; the coastal subspecies migrates south along the North American Pacific Coast to Mexico and Central America. The inland subspecies migrates southeast, through the eastern United States to South America^[27]^. Ecological niche modeling in this system indicates parental populations occupy different ecological niches on the breeding grounds and migration^[28,29]^. Niche modeling has not been extended to the wintering grounds but these differences likely extend to the wintering grounds where they occupy geographically distinct areas^[25]^. Following Rice and McQuillan^[30]^, these ecological differences could have led to divergent selection on learning abilities and reduce the fitness of hybrids. Paleodistribution models also suggest coastal and inland thrushes were isolated in different refugia during the last glacial maximum (i.e., they were allopatric)^[28]^. Alleles underlying cognitive traits could have diverged during this period and be incompatible in present day hybrids, causing misexpression at neurological mechanisms that support cognition. Independent of cognition, misexpression can also increase cellular stress and have several downstream effects including reduced cognitive performance^[31]^.

The primary objective of the present study was to compare the relative learning and memory abilities of parental and hybrid thrushes. We used two cognitive tests for this work – an associative learning spatial test and neophobia test. These tests assess domain-general aspects of learning and memory that are important for myriad foraging tasks and managing unpredictable environments^[32,33]^. Although these specific tests are most commonly applied to food-caching birds, the taxa on which the bulk of study on foraging-related memory and learning occur in birds^[16,17,33]^, they assess the accuracy of basic cognitive skills required for any foraging task – memory and problem solving. Consequently, differences in performance on these tasks should represent essential, albeit simplified (i.e., measured in a laboratory-based setting) differences in cognitive abilities among taxonomic groups that scale up to ecologically-relevant phenomena.

Thus, we predicted that if learning and memory serve as post-mating isolating barriers in the Swainson’s thrush, hybrids would perform less well than parental forms on these tests. We had two additional objectives in the present study as well, to test the effects of sex and season on learning and memory. Specifically, in their study of hybrid chickadees, McQuillan et al. 2018^[16]^ noted that female hybrids performed worse than male hybrids. This finding supports Haldane’s Rule, that the heterogametic sex (females in birds) experiences greater fitness losses. We set out to test this hypothesis in thrushes as well, comparing the cognitive abilities of male and female thrushes. We also ran our experiment during both the non-migratory and migratory season. Because cognitive demands are likely higher during migration, when birds are developing navigational maps for subsequence migrations and as they enter new environments at stopover sites, we predicted that birds (regardless of sex or their status as parental or hybrid forms) would perform better in cognitive tests during the migratory season if cognitive demands are higher during migration.

## METHODS

All experiments were performed in accordance with relevant guidelines and regulations. Protocols were approved by the Institutional Animal Care and Use Committee at Texas A&M (IACUC 2019-0066) and permits were obtained from Environment and Climate Change Canada (SC-BC-2020-0016), the U.S. Fish and Wildlife Service (MB49986D-0) and Texas Parks and Wildlife Commission (SPR-0419-067). The study is reported in accordance with ARRIVE guidelines.

### (a) Collection sites

We captured juvenile birds using mistnets and song playback at the end of the breeding season (August 2021) in British Columbia, Canada and brought them to our vivarium in College Station, Texas. Fifty-eight birds in total were used; 19 birds from the coastal range (Vancouver; 10 males and nine females), 20 birds from the hybrid zone (Pemberton; 10 males and 10 females) and 19 birds from the inland range (Kamloops; 10 males and nine females^[26]^).

Songbirds become nocturnal when they migrate and exhibit migratory restlessness (*zugunruhe*). To monitor this transition and ensure we were testing the thrushes at appropriate times, we used motion sensors designed at Texas A&M University^[34]^. Each cage was equipped with a AM312 Passive Infrared (PIR) sensor (HiLetGo, Guangdong, China) that detected any movement of the individual. All PIR sensors were powered by a protoboard and wired to a central unit using a Data Acquisition Card (DAQ). The DAQ interfaced the motion detections to the LabVIEW program on a central Windows Operating System computer. Data was output into a CSV file in 10-minute increments. We validated the reliability of motion sensors by collecting behavioral data from a subset of birds (n=21) using infrared (IR) cameras (*D-link, DCS-932L Day/Night Network Surveillance Camera*). We found that data from PIR sensors and IR cameras were highly correlated^[34]^.

We classified birds as non-migratory and migratory based on similar criteria to Johnston et al.^[35]^. The PIR sensors have a three-second delay after the detection of motion, so we divided motion detection records by three to estimate a more accurate movement count. We defined time increments as active when a bird moved greater than 20 times per 10 minutes. The number of increments per day was adjusted according to the photoperiod. Non-migratory birds were active for less than 5% of nightly time increments and migratory birds were active for greater than 40% of nightly time increments.

We ran experiments during both winter (non-migratory, Jan 2022) and spring migration (Mar 2022). All birds were run through cognitive tests during the winter. A subset of birds was sacrificed for another experiment leaving 29 birds for the spring season (5 males and 5 females from Vancouver; 4 males and 5 females from Pemberton; and 5 males and 4 females from Kamloops).

### (b) Housing

We housed birds individually and fed them a diet consisting of berries, mealworms, egg, crackers, cottage cheese and red meat *ad libitum*^[35]^. Photoperiod was gradually decreased in September-January from 12.75 hours of light (L):11.25 hours of dark (D) to 11L:13D. We used geolocator data from hybrid thrushes to estimate timing and pace of migration and adjust timing of photoperiodic changes^[27,36]^ Birds were held at 11L:13D until February. In February–March, photoperiod was then gradually increased to 16L:8D to mimic early spring on the breeding grounds. All behavioral tests were performed in their home cages (24” × 13” × 12”).

### (c) Learning tests

#### (i) Associative learning

We followed methods described in Roth et al. (2012) for this task^[33]^, testing how many trials it took for birds to find two worms hidden in a particle board memory board (∼30 × 30 cm square board; ∼1.25 cm diameter holes; Figure S1) with 16 wells. Black pom-poms were inserted into each well, which the birds had to remove to access a mealworm treat. We started with a period of acclimation, gradually introducing the birds to particle boards over ten days. Pompoms were replaced every day; birds were considered acclimated when they removed all of the pom-poms for two consecutive days.

To control for motivation changes that might be affected by hunger, the birds were given access to their usual diet for up to 2 hours before any tasks. For the main trial, we placed two mealworms in a single well, covered all the wells with pom-poms and ran the trial for three hours for ten consecutive days (using the same well each day and starting trials at 11 AM). We recorded the number of pompoms removed until the correct well was located. This experiment was performed twice, once in December-January (migratory period) with the whole cohort (n=58), and again in March with the remaining half (n=29) that were not sacrificed in January.

Wells containing worms were different for each season and chosen at random. Accordingly, we think it unlikely that well position affected our results (e.g., all hybrids were not assigned wells on the inside of the board). In support of this suggestion, there was no relationship between ancestry group and well position (“inside” or “outside” the board; χ^2^ = 8.23, p = 0.09). In addition, we did not document any difference in performance based on well position (p = 0.12 in generalized linear model run as described below under ‘Statistical Analysis’).

Beyond well position, differences in motivation could also have affected our results. We tested for this effect in several ways. First, we performed pre- and post-trial controls, placing a mealworm on top of the boards, and quantifying how long it took for the bird to land and take the worm. No differences were documented between the groups (ancestry or sex) or seasons (all p > 0.06 in linear model run with ancestry, sex and season as predictor variables and individual as a random effect). Second, beyond pre- and post-trial controls, we also documented the time it took birds to land on the boards each day of the actual trial and found no differences between the groups or seasons (all p > 0.083 in linear models run as described below under ‘Statistical Analyses’ but with seconds instead of score as the response variable). Third, the first day of the actual trial was not included in our analysis as we would expect birds to perform randomly that day. No differences were documented between the groups or seasons in how well they performed that day (all p > 0.34 in generalized linear models run as described below under ‘Statistical Analyses’). This may be a better control than testing how long it took them to take mealworms from the top of the board or land on boards because it required birds to search for their food. Finally, we also compared average scaled mass indices across groups and seasons^[37]^. These indices were calculated using body masses measured before the trial in winter and spring and scaled by tarsus length. The only difference we documented was by season, with birds having higher indices in the winter (p < 0.001; all other p > 0.l8 in linear model run with ancestry, sex and season as predictor variables and individual as a random effect). If increased body mass reduced motivation, we would expect birds to perform worse in the winter. That is not the case (see ‘Results’) and thus we do not think difference in body mass affected our final results.

#### (ii) Response to novelty

Following the associative learning trials, we performed a novel latency trail. Feeding conditions were kept the same as the associative learning task, allowing the birds to feed for up to 2 hours before the trial. For this task, we created a novel object by modifying the birds’ normal white feed bowl. During winter, the bowls were painted red and binder clips were added to all sides; during the spring, bowls were painted blue and paper clips were added to all sides (Figure S1). In the trial, four meal worms were placed into the novel feed bowls and the trial was run for one hour starting at 9AM. Birds were randomly split into two groups for this trial so it could be run over two days, allowing food to be put in quickly with limited disturbance. We recorded the times at which the bird flew down to inspect the novel bowl, the time they ate the worm and the total number of worms eaten. Similar to the associative learning protocol, a pretrial and post-trial control was executed the same day to control for motivation. For the controls, the birds were given their familiar feed bowls with four mealworms 10 minutes before and after the novel trial.

### (d) Statistical analyses

#### (i) Associative learning

We limited this analysis to birds that were assayed in both winter and spring migration (n = 29) and fit generalized linear-mixed models (GLMMs) with log link functions in the “lme4” package of R to our data. We included “score” (the number of pom-pom balls removed before the bird recovered the mealworms) as the response variable in these analyses and four fixed effects (along with all possible interactions): ancestry (coastal, hybrid or inland), sex, testing day, and season (winter or spring migration). We specified a random intercept and slope for each bird across testing days and used step-wise model simplification, removing the least significant variable (starting with the highest order interactions) until we reached our best-fit model. Likelihood ratio tests (LRT) were used in each step, comparing models with and without focal terms. If the simplified model explained significantly less variation in the response variable the focal term was retained. Post-hoc tests were conducted using least-square means (LSM) in the R package “lsmeans”. The first day of this trial was not included in the analysis as birds would be expected to perform randomly that day.

#### (ii) Response to novelty

We ran separate analyses for each season as different novel feeders were used precluding a comparison across seasons (n = 58 for winter and 29 for spring). Repeated-measures analyses of variance were used (using the aov function in R), testing for a between-subjects effect of population and within-subject effect of control vs. treatment. Separate analyses were run for latency to approach and eat from the feeders. Post-hoc tests were conducted using general linear hypotheses (GLHT) in the R package “multcomp”. Data were log-transformed for all analyses but raw data are presented in figures for clarity.

Ancestry was included as a categorical variable (hybrid, coastal or inland) in all of our analyses. Roughly 40% of the birds in Pemberton are hybrids^[26]^. We genotyped birds in the field using three RFLPs diagnostic of inland and coastal subspecies^[36]^ and kept birds with the highest degree of admixture. Ancestry could range from 0 (coastal, all three RFLPs are homozygous for coastal alleles) to 1 (inland, all three RFLPs are homozygous for inland alleles). Average ancestry of birds included in experiment was 0.50 (range 0.17-0.83). Males were subsequently sequenced with a whole genome resequencing approach and ancestry estimated as described in Delmore et al. 2016^[38]^. Results from analyses remained the same when including ancestry as a continuous variable (e.g., Table S1 and Figure S2).

## RESULTS

### (a) Associative learning

Associative learning was quantified as the number of pompoms removed before the food reward was found. Males and females were tested, and the same individuals were assayed during the winter (non-migratory) and during spring migration. All groups showed evidence of learning; none of the groups performed better than random expectations (eight pompoms removed, or “inspections”) on the first day of the experiment (one-sample t-tests, all p > 0.05) but all groups fell below random by the end of the experiment (one-sample t-tests, all p < 0.0001). We focused the remaining analyses on results from days 2-10 (i.e., after birds were introduced to the task and had the chance to learn).

The best-fit model included all fixed effects and interaction terms (LRT compared to null model: χ^2^(23) = 83.83, p < 0.0001; LRT comparing models with and without highest order interaction [ancestry:sex:day:season]: χ^2^ (2) = 9.89, p = 0.0071; Table 1; Figure 1). As suggested by the former one-sample t-tests, testing day was a significant negative predictor of performance, indicating learning occurred across all ancestry groups (i.e., all birds required fewer inspections to recover the food reward as the test progressed, χ^2^ = 57.94, p < 0.0001). Season was also a significant predictor of performance but, contrary to our predictions, fewer inspections were required to recover the food reward during the winter (vs. spring migration) for all ancestry groups (χ^2^ = 9.14, p = 0.0025). Significant χ^2^ values for interaction terms sex:season (χ^2^ = 8.64, p = 0.0032) and ancestry:sex:season (χ^2^ = 6.12, p = 0.047) indicate that this decrease in the number of inspections required during the winter was especially true for inland males. Of prime importance for our study, the highest order interaction term in our best fit model (ancestry:sex:day:season) was significant (χ^2^ = 10.00, p = 0.0067). In line with our prediction that hybrid thrushes would exhibit reduced cognitive performance compared to parental thrushes, hybrid males required more inspections to recover food than males from either parental population at the beginning of the experiment. This pattern was only true during the winter (Figure 1 top right panel).

**>Table 1.** Results from GLMM examining the relationship between score (the performance of birds at an associative learning spatial task) and a series of predictor variables.

|  | X<sup>2</sup> | df | pvalue |
| --- | --- | --- | --- |
| ancestry | 0.64 | 2 | 0.72 |
| sex | 0.00 | 1 | 0.96 |
| **test_day** | **57.94** | **1** | **<0.0001** |
| **season** | **9.14** | **1** | **0.0025** |
| ancestry:sex | 3.70 | 2 | 0.16 |
| ancestry:test_day | 1.81 | 2 | 0.41 |
| sex:test_day | 0.81 | 1 | 0.37 |
| ancestry:season | 5.26 | 2 | 0.072 |
| **sex:season** | **8.64** | **1** | **0.0033** |
| test_day:season | 1.87 | 1 | 0.17 |
| ancestry:sex:test_day | 3.99 | 2 | 0.14 |
| **ancestry:sex:season** | **6.12** | **2** | **0.047** |
| ancestry:test_day:season | 3.55 | 2 | 0.17 |
| sex:test_day:season | 0.22 | 1 | 0.64 |
| **ancestry:sex:test_day:season** | **10.00** | **2** | **0.0067** |

**Figure 1.**
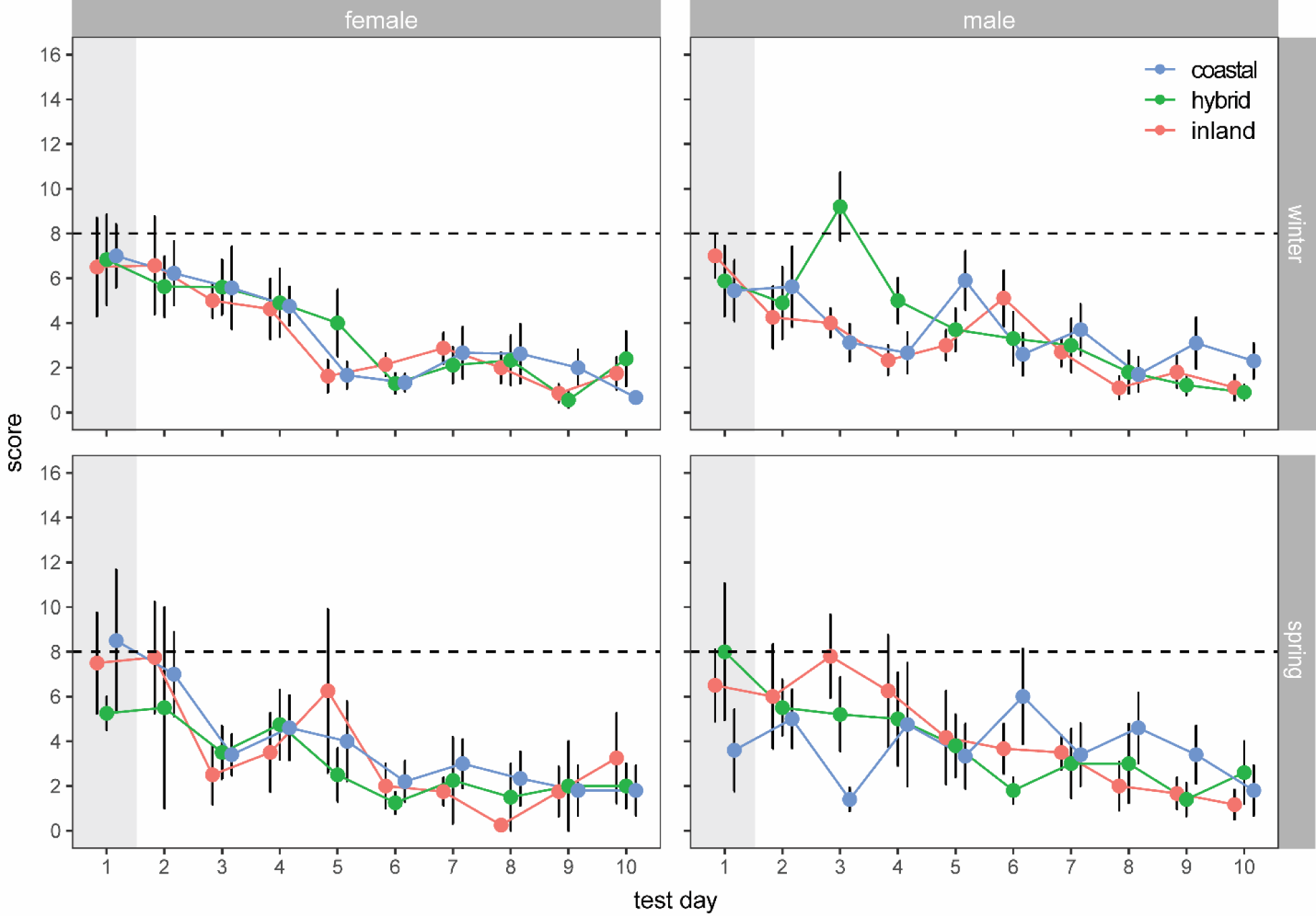
Results from associative learning spatial task, showing performance over the testing period for all three ancestry groups in separate seasons for each sex. Score refers to the number of inspections required to obtain the food reward. Top right panel shows hybrid males required more inspections than parental forms during the winter (χ^2^ = 10.00, p = 0.0067). Day 1 is grayed out because it was not included in our formal analyses; birds are expected to perform randomly that day (one-sample t-tests validate this suggestion, see text). Means (+/- standard errors) are shown.

Before moving on to the response to novelty experiment, we took a closer look at results for males from winter. First, we used one-sample t-tests to compare the mean number of inspections required by each ancestry class to random expectations (eight inspections). As noted already, none of the ancestry groups performed better than random on the first day. As expected based on results from our larger model, pure forms fell below random expectations before hybrids, with inland birds requiring fewer inspections than random by day two (mean score = 4.25, p = 0.008), coastal birds by day three (mean score = 3.16, p = 0.006), and hybrids by day five (mean score = 3.7, p = 0.001). Note, hybrid birds required an especially large number of inspections on day three. We reran our larger model randomly replacing hybrid scores from day three with scores from days one and two and the results remained the same (hybrids still required more inspections early in the experiment) suggesting results from day three are not driving the patterns we documented. In addition, as noted above with one-sample t-tests, hybrids did not fall below random expectations until day five (i.e., their performance was also low on days two and four of the experiment).

### (b) Response to novelty

Response to novelty was quantified as the latency to both approach and eat from a novel feeder. We controlled for differences in motivation by quantifying these latency variables pre-trial and post-trial with their normal feeder. Separate models were run for winter and spring migration. Birds responded to the novel feeder in all cases, taking longer to approach the novel vs. normal feeders (all p < 0.0001; comparing pre and post to treatment in Figure 2 and S3). Population was a significant predictor of the latency to approach and eat from the feeder during the winter (approach: F_2,50_ = 4.30, p = 0.02; eat: F_2,50_ = 3.39, p = 0.042; Figure 2 and S3) but the pattern was not as predicted; inland birds (not hybrids) exhibited greater latency than both hybrids and coastal birds (approach – comparison to hybrids: estimate = 0.21, p = 0.11, comparison to coastal: estimate = 0.29, p < 0.02; eat – comparison to hybrids: estimate = 0.21, p =0.064, comparison to coastal: estimate = 0.021, p =0.09). Ancestry was not a significant predictor of latency to approach or eat from the feeder during spring migration (approach: F_2,23_ = 0.93, p = 0.41; eat: F_2,23_ = 0.49, p = 0.62).

**Figure 2.**
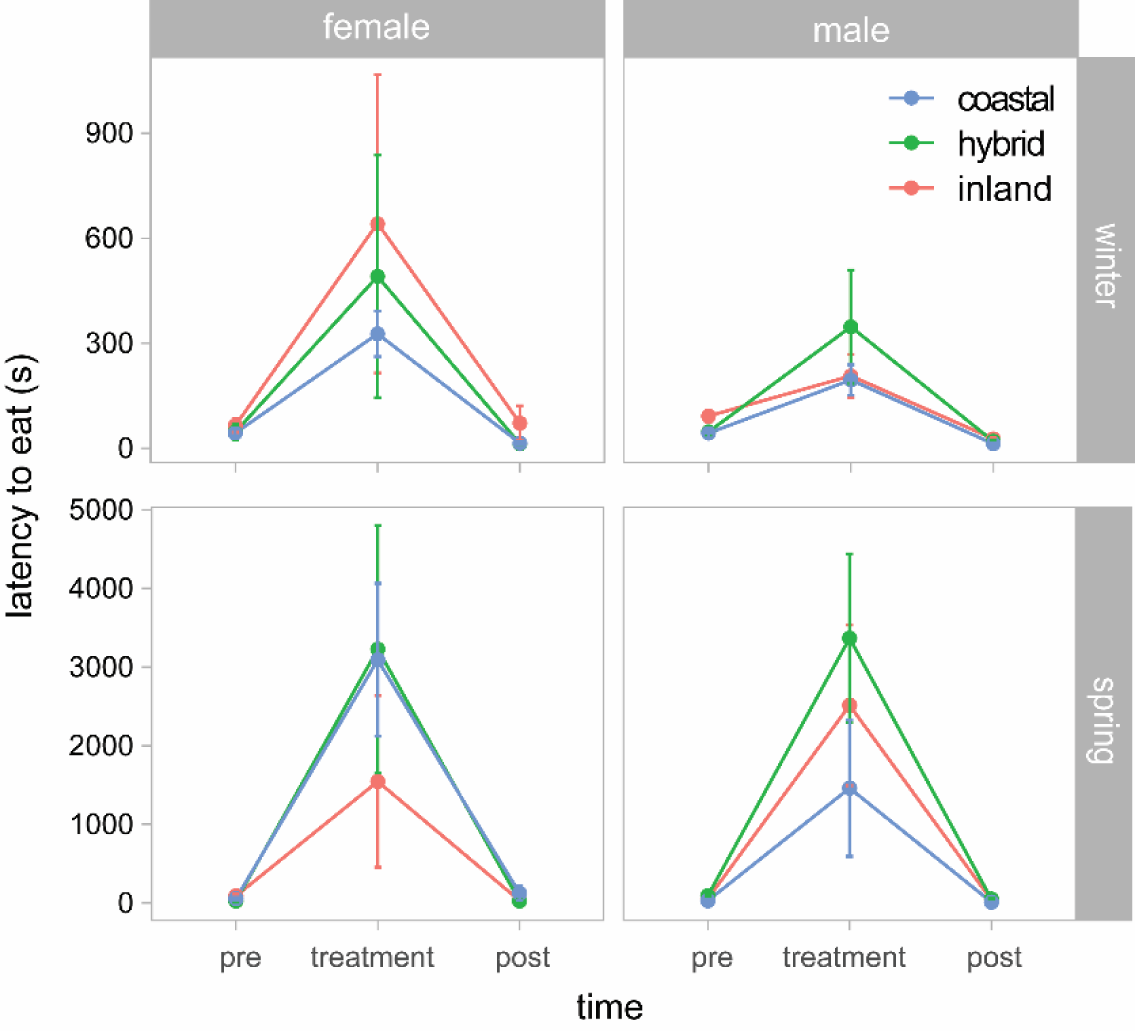
Results from response to novelty task, showing latency to eat from a feeder. Normal feeder used pre- and post-treatment. Novel feeder used for treatment. Mean +/- standard error shown. Analyses were run using logged values for latency to eat but untransformed values are shown here for clarity. Results for latency to approach feeder shown in Figure S3. Birds took significantly longer to eat from novel dishes (treatment) in all cases; hybrids did not require significantly more time to eat in any comparison.

## DISCUSSION

We assayed two general cognitive factors (learning about a novel object and a spatial memory task) in a hybrid zone between Swainson’s thrushes. We predicted that hybrids would perform worse than parental forms if these traits serve as post-mating isolating barriers. We assayed the cognitive abilities of both males and females and ran our experiments during both winter and spring migration, predicting that hybrid females (the heterogametic sex) would perform worse than males and birds (regardless of ancestry or sex) would perform better on spring migration. We found that ancestry alone was not a significant predictor of performance in the spatial memory task, nor was the interaction term between ancestry and sex (i.e., hybrid females did not perform worse). We did find some indication that ancestry affects spatial memory performance, but this was limited to wintering hybrid males. Although season was a significant predictor of spatial memory performance, contrary to our predictions, birds learned faster during the winter relative to spring migration. In contrast, ancestry was a significant predictor of novel learning success, although inland birds actually performed worse than both hybrids and coastal birds, without an effect of sexes or season.

### Cognition as a post-mating reproductive isolating barrier

Rice and McQuillan^[30]^ described two ways learning and memory could serve as post-mating isolating barriers, with hybrids between ecologically divergent parental populations exhibiting intermediate and inferior cognitive abilities and/or alleles underlying cognition fixing during periods of allopatry and causing misexpression when they are recombined in hybrids. Both scenarios could apply to Swainson’s thrushes, as they occupy different ecological niches^[28,29]^, and were likely isolated in different refugia during the glacial cycles of the Pleistocene^[28]^. In line with this suggestion, we found some support for learning and memory as post-mating isolating barriers here. Although ancestry alone did not predict performance in either of the cognitive tasks we examined, we did document a reduction in performance by male hybrids during the winter in the associative learning spatial task. Evidence from a complementary study using these birds suggests misexpression may play an important role in reducing hybrid male cognition. Specifically, transcriptomic analyses using these birds showed considerable misexpression in the hippocampus of these individuals. This pattern was limited to the winter (vs. spring) and a comparable increase in misexpression during the winter was not documented in the other brain regions examined (Cluster N and the hypothalamus; unpublished data).

Reductions in learning and memory could affect any number of fitness-related traits in thrushes. The tests we used are related to resource acquisition, e.g., foraging and managing unpredictable environments^[32,33]^. Swainson’s thrushes do not form pairs on the wintering grounds^[39]^, but difficulties acquiring resources on the wintering grounds could carry-over to spring migration and the breeding season, affecting hybrid survival on migration and reproductive success on the breeding grounds^[40,41]^. These effects on hybrid fitness could reduce gene flow between Swainson’s thrushes and contribute to reproductive isolation in the system. It is important to note that the cognitive deficits we documented here could derive from reductions in hybrid fitness that are related to larger scale physiological deficiencies such as general reductions in health or body condition that affect factors such as motivation. In addition, it is difficult to say how effective this barrier to gene flow is in Swainson’s thrushes, as it only affects one sex and season. Very few studies have compared the cognitive abilities of parental and hybrid forms. McQuillan et al. (2018)^[16]^ documented a greater effect of ancestry on the cognitive performance of hybrid chickadees during the fall/winter, but they did not include a comparison across seasons.

The relationship we documented between ancestry and sex is not consistent with our predictions. Haldane’s rule predicts that the heterogametic sex (females, in birds) will exhibit lower fitness than the homogametic sex. However, in this study, males exhibited significantly lower cognitive abilities. This contradictory finding may reflect the fact that Swainson’s thrushes have not been diverging for very long; mitochondrial divergence between the subspecies is 0.69^[25]^. Combined with a molecular clock of 2% divergence per million years, this estimate of divergence suggests Swainson’s thrushes began diverging 350 thousand years ago. Haldane’s rule derives from the accumulation of genetic incompatibilities between parental genomes. These incompatibilities can take a long time to accumulate, especially in birds where it can take millions of years for this form of intrinsic, postzygotic isolation to evolve^[42–44]^.

We look forward to future work interrogating the relationship we documented between cognitive performance and ancestry, season and sex. For example, future work using a larger sample size of birds would be valuable, as it could allow us to both validate our results and allow for more detailed analyses (e.g., examining changes in performance each day rather than over the full testing period, ensuring one or two days are not driving our results). In addition, it is possible that birds used olfactory cues (instead of cognition) to find worms. Probe trials could be used to test for this effect, quantifying performance at the end the trial when no food is available anywhere in the board. If birds are using olfactory cues their performance should be random during this trial.

### Potential differences in cognition between the seasons

We expected memory differences between seasons to be apparent during the migratory period. Large-scale navigation during migration is typically associated with enhanced memory capacity in general and enhanced resources devoted to neurological mechanisms that enhance memory during the migratory period. Although birds, including Swainson’s thrushes, undoubtedly use multiple navigational mechanisms during migration such as geomagnetic, celestial, and olfactory cues (reviewed in ^[45]^), evidence strongly suggests that experience, spatial learning, and cognition are also important components of avian migration, especially in terms of landmark-based navigation and the creation of experience-based, large-scale cognitive maps^[45–51]^.

Although we expected that season was a significant predictor of spatial memory performance, we found instead the opposite - enhanced memory performance occurring during the winter. Most migratory cognition predictions focus on the fall – gearing up for and getting to the wintering grounds^[45]^. In our study, we were specifically focused on the periods of activity on the wintering grounds and during migration back to the breeding grounds. If cognition is important for migration in this species, then we might actually predict that cognitive processes might be less important on the return trip to the breeding ground. In other words, if the bulk of the learning had already occurred during the fall, then the return to the breeding grounds might be less cognitively demanding, as the birds would simply be repeating it, although in reverse. As such, we might see such a reduction in cognitive activity.

Likewise, it is possible that Swainsons’ thrushes do not depend greatly on cognitive processes during migration and instead use other strategies during migration. Swainson’s thrushes might rely on less cognitively intensive cues for migration, such as those facilitated by cryptochromes for magnetic field orientation with twilight cues to calibrate the magnetic compass^[52,53]^. In this case, we might not expect thrushes to perform better during migration (spring or fall), but instead during the wintering period. Consistent with this idea, de Morais Magalhães et al. ^[54]^ suggest that semipalmated sandpipers (*Calidris pusilla*) have high levels of hippocampal neurogenesis in wintering birds, perhaps as a function of recovery from or a response to migration. The physiological consequence of exercise^[55]^ or stress during migration^[56]^ might warrant an increase in neuron production and maintenance after migration, which might affect behavioral patterns on the wintering site. Although speculative, we note that enhanced cognitive performance on the wintering site might be adaptive as it allows for enhanced resource acquisition, preparation^[57]^, and thus earlier departure from the wintering grounds, earlier arrival at the breeding grounds, and so possibly better access to breeding territories^[58]^. Nevertheless, we cannot dissociate these effects from those of changes in motivation and the need to divert resources and attention to non-cognitive migratory factors (e.g., gearing up for reproduction, increasing HVC for song production^[59]^) that might occur independently of spatial or other cognitive processes during the spring migration). Future work (e.g., assaying hormones like testosterone and corticosterone that may vary by season) will help dissociate these effects. And we note that birds assayed in spring were in captivity for longer. There is some evidence the hippocampus is negatively affected by captivity^[60]^ so future work controlling for time in captivity is also needed.

### Conclusion

Although we found evidence that seasonality and to a lesser extend affect the cognitive abilities of migratory Swainson’s thrushes, the patterns we documented contradicted several of our predictions that were based primarily on food-caching species (the most well studied paradigm for comparisons across avian hybrid cognition and behavior^[16]^) and more general patterns of cognitive performance with changing migratory status^[45]^. There may be a trend towards hybrid males performing worse in novelty assays (Figure 2; Figure S3); future work using larger sample sizes may support this trend. Additional behavioral assays may also uncover differences in a cognitive traits we did not assay here. A key next step will also be to examine these phenomena in the context of changes in the mechanisms that support variation in cognitive capacity, e.g., neural structures, particularly as they relate to sex differences and season.

Work with the hippocampus supports a role for learning during migration. The hippocampus is important for learning and spatial orientation^[61,62]^. As such, migratory species tend to show enlarged hippocampal formation volume when compared to non-migratory species^[63–67]^. Moreover, the hippocampus can respond with age and experience that is associated with higher spatial memory demands^[68]^, resulting in seasonal changes even in species that do not migrate^[64,68]^. To the best of our knowledge, no one has compared the relationship between learning and memory and the associated neural correlates between the migratory and non-migratory season in a single species across a hybrid zone or across various seasons of migration. The neural correlates of learning about novel stimuli are less well known, although may be associated with the arcopallium (the amygdala homolog in birds^[33]^). Given the behavioral results herein, we would predict enhanced hippocampal volume and/or neuron density in males, and a shift in hippocampal size in preparation for the fall migration, lasting into the winter. This difference should be reduced in the spring. We would also predict inland birds to possess larger arcopallium regions that are maintained throughout the year.

## Supporting information

Supp Mat

## Acknowledgements

This research was supported by grants from NSF to KED (IOS-2143004) and TCR (IOS-1754297). Also a mini research grant to AA from the Schubot Center for Avian Health. We thank members of the Delmore lab for assistance during field work and the captive experiment; Scott Taylor and Amber Rice for reading drafts of the manuscript.

## Author contributions

TCR and KED designed the experiment; AA, HJ, CW and KED carried out the experiment; KED performed analyses; AA, MT, TCR and KED wrote manuscript with input from HJ and CW.

## Data availability

All data generated or analysed during this study are included as electronic supplementary material, including data to run associative learning (al.csv) and neophobia (neophobia_*.csv) analyses.

