## Supplementary material for "Learning and memory in hybrid migratory songbirds - cognition as a reproductive isolating barrier across seasons": Supp Mat

**Table S1.** Results from GLMM examining the relationship between score (the performance of birds at an associative learning spatial task) and a series of predictor variables. Ancestry coded as continuous variable in these analyses (vs. categorical as in Table 1).

|  | X^2^ | df | p-value |
| --- | --- | --- | --- |
| ancestry  test_day  season  ancestry:test_day  ancestry:season  test_day:season  ancestry:test_day:season | 3.39  34.16  17.40  1.46  13.24  0.50  4.51 | 1  1  1  1  1  1  1 | 0.065  **<0.0001**  **<0.0001**  0.23  **0.00027**  0.48  **0.034** |

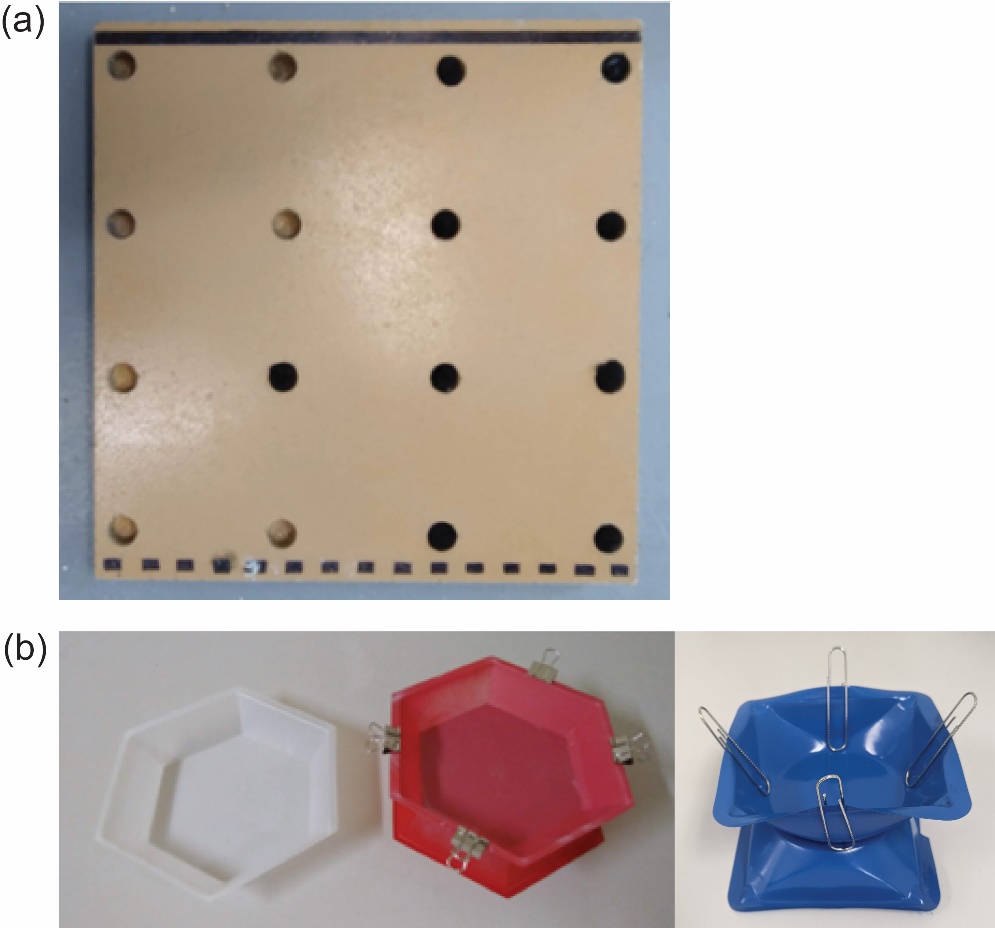

**Figure S1**. (a) Board used for associative learning spatial test and (b) food dishes used for novel problem-solving test (white = normal dish, red = novel dish for winter, blue = novel dish for spring migration).

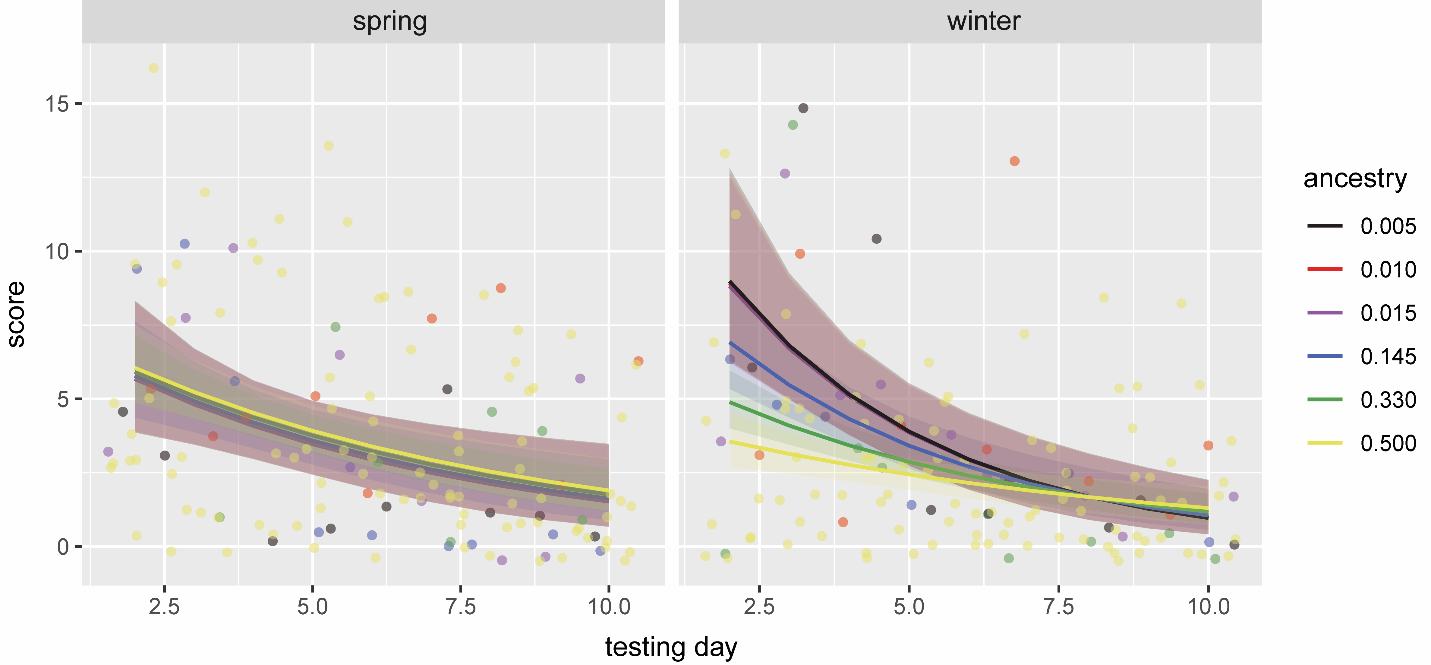

**Figure S2**. Results from associative learning spatial task, showing performance over the testing period for all three ancestry groups in separate seasons. Score refers to the number of inspections required to obtain the food reward. Ancestry coded as continuous variable with a value of zero indicating F1 hybrids and 0.5 pure coastal or inland thrushes (vs. categorical as in Figure 1). Right panel shows hybrid males required more inspections than parental forms during the winter.

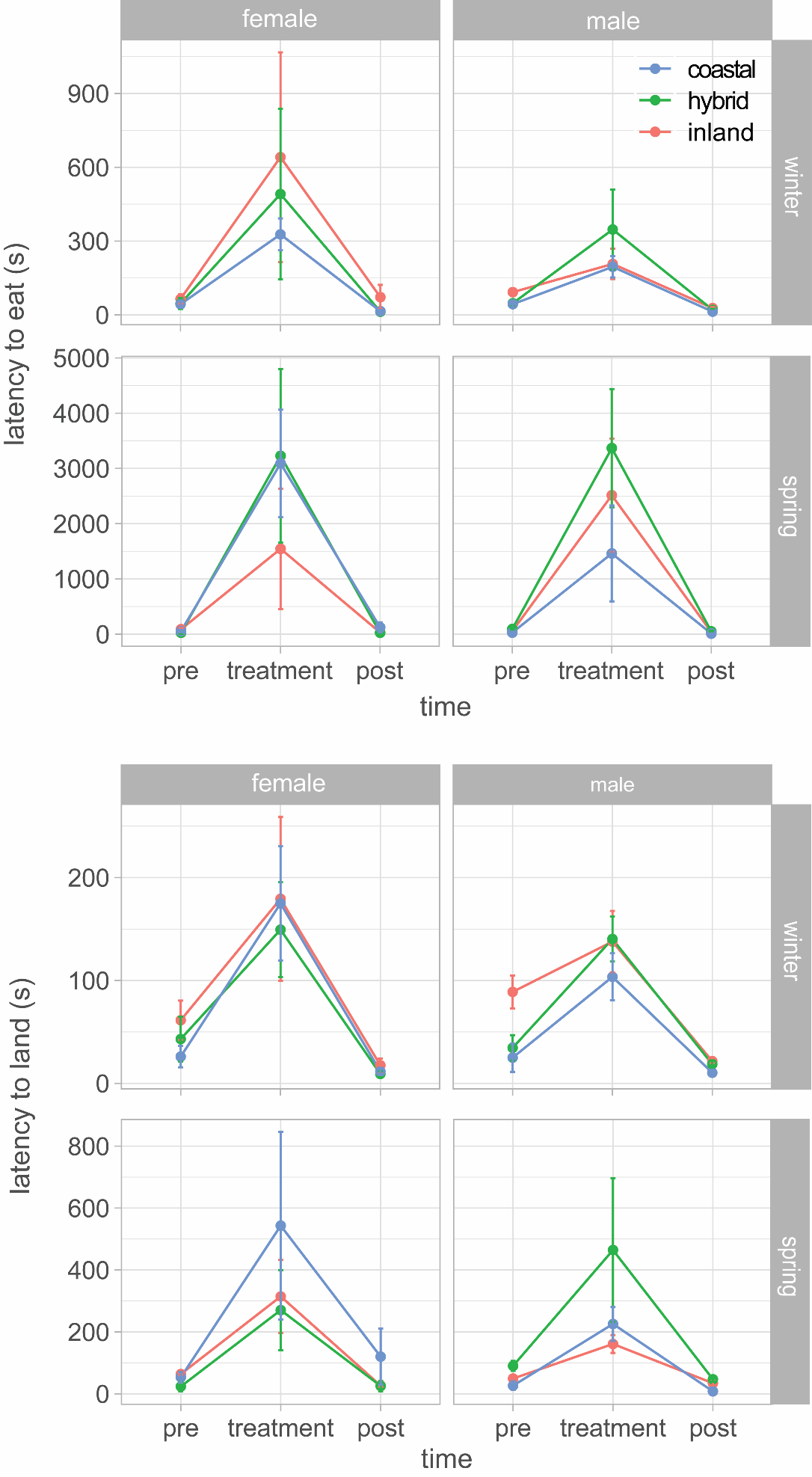

**Figure S3**. Results from response to novelty task, showing latency to approach feeder. Normal feeder used pre- and post-treatment. Novel feeder used for treatment. Mean +/- SE shown. Analyses were run using logged values for latency to eat but untransformed values are shown here for clarity. Results for latency to eat from feeder shown in Figure 2. Hybrids did not require more time to eat in any comparison. Blue = coastal; Green = hybrid; Pink = inland.
